# The role of miscarriage and sororal birth order in male same-sex orientation: Theoretical predictions and empirical data

**DOI:** 10.64898/2026.03.09.710348

**Authors:** Michel Raymond, Ana Aguerre, Valerie Durand, Menelaos Apostolou, Julien Barthes, Sarah Nila, Bambang Suryobroto, Mostafa Sadr-Bazzaz, Paul L. Vasey, Daniel Turek, Pierre-André Crochet

## Abstract

This study explores the proximal and biological mechanisms underlying male same-sex orientation, with a focus on the Fraternal Birth Order Effect (FBOE), a robust phenomenon whereby same-sex oriented men tend to have more older brothers than the population average, and its relationship with the Sororal Birth Order Effect (SBOE), whereby older sisters also appear to influence sexual orientation, albeit less consistently. The Maternal Immune Hypothesis (MIH), which posits that maternal immune responses to male-specific antigens accumulate across successive male pregnancies, provides a compelling proximal explanation for the FBOE, but it fails to fully account for the SBOE and other birth order patterns, such as the elevated prevalence of same-sex orientation among only-children compared to firstborns in larger sibships. Through explicit modelling of the MIH, our simulations reveal that the correlation between the number of older brothers and sisters generates a spurious SBOE, which disappears when controlling for older brothers. However, this control becomes insufficient when miscarriages are included in the simulations. Additionally, the increased prevalence of same-sex orientation among only-children, relative to firstborns with siblings, only emerges when miscarriages are incorporated into the model. Empirical analyses across eight diverse populations (Indonesia, France, French Polynesia, Greece, Canada, Czech Republic, Samoa, Iran) confirm the presence of an overall significant FBOE and, critically, an overall significant SBOE even after controlling for the number of older brothers. The higher frequency of same-sex orientation men among only-children, compared to firstborns in larger sibships, supports a possible role of miscarriage for this SBOE. However, the estimated miscarriage rates (37% - 58%) explaining the observed SBOE slightly exceeds reported rates (30% - 45%) which conservatively include early pregnancy losses. This possibly suggests that additional mechanisms contribute to a spurious SBOE or that a genuine SBOE coexists with the FBOE. Limitations of this study are discussed, as well as whether the MIH framework can be extended to accommodate these findings, or if alternative explanations are needed to resolve these discrepancies.

## Introduction

Male same-sex orientation toward physically mature members of their own sex (i.e., male androphilia), for sexual intercourse, romantic relationships, or both, is an evolutionary enigma. This is because preference for male-male relationships is partially heritable (Bailey et al., 2000; Långström et al., 2010) and is associated with a fertility cost with a 30-100% decrease in offspring number (Iemmola & Camperio-Ciani, 2009; Nila et al., 2018; Rieger et al., 2012; Vasey et al., 2014). Also, male androphilia is surprisingly common in many societies (2%–6% in Western countries) for such a costly trait (Apostolou, 2020b; Berman, 2003; Rahman et al., 2020).

A solid empirical observation is that androphilic males have on average more older brothers compared to heterosexual men: this is referred to as the fraternal birth order effect, or FBOE (Blanchard & Bogaert, 1996; Blanchard & Klassen, 1997). The FBOE has been found independently in Western (e.g., Ablaza et al., 2022; Apostolou, 2020a; Blanchard, 2018b, 2018a; Blanchard & Bogaert, 2004; Bogaert & Skorska, 2011; Fořt et al., 2024) and non-Western countries such as Turkey, Iran, Hong Kong, Samoa, Mexico, and Indonesia (Blanchard, 2018a; Gómez Jiménez et al., 2020; Li & Wong, 2018; Nila et al., 2019; Sadr-Bazzaz & Vasey, 2025; Semenyna et al., 2023). The only exceptions to this empirical rule appear to come from studies with small sample sizes or populations where the average fertility rate is low, e.g., France (Raymond et al., 2023), China (Xu & Zheng, 2017), Thailand (Skorska et al., 2020). The FBOE thus remains a well-established proximal determinant of male androphilia.

The FBOE has not revealed any statistical association with potential confounders such as sibling size, birth year, age, socio-economic status (Blanchard & Bogaert, 1996; Bogaert, 2003) or parental age (e.g., Bogaert, 2006; Bogaert & Liu, 2006). Most importantly, it seems to be present even when the biological brothers were raised in different households, suggesting that it operates only during prenatal life (Bogaert, 2006). The FBOE thus seems to be caused by lasting effects from previous male gestations. The first proposed candidate for such “memory” across the successive pregnancies has been the maternal immune system: under the maternal immune hypothesis (MIH), mothers would develop antibodies against proteins involved in male brain development such as Y-linked proteins, with increasingly stronger effects with each male pregnancy, affecting the brain structures underlying sexual partner preference in later-born sons (Blanchard et al., 2001; Bogaert & Skorska, 2011). A recent study provided empirical support to this hypothesis by showing that mothers of gay sons with older brothers had significantly higher circulating levels of anti-neuroligin 4 Y-linked (NLGN4Y, a Y-linked protein) antibodies than the control samples including mothers of only gynephilic (i.e., attracted to physically mature women) sons (Bogaert et al., 2018). In spite of this apparent empirical support for the MIH, three lines of evidence are challenging the validity of this hypothesis to explain the FBOE. Recently, three additional birth order effects on sexual orientation have been identified, suggesting that the MIH is either inadequate or needs to be supplemented with other hypotheses to proximally explain all birth order effects on sexual orientation.

First, a sororal birth order effect (SBOE) has been described, which is not predicted by MIH. It has been described in several studies, e.g., in UK (King et al., 2005), Finland (Kangassalo et al., 2011), Samoa (Semenyna et al., 2023), Canada (Swift-Gallant et al., 2018), Iran (Sadr-Bazzaz & Vasey, 2025), and confirmed in a meta-analysis (Blanchard et al., 2021) and in a nation-wide study (Ablaza et al., 2022). This SBOE has always been found in association with a FBOE and has usually been found to be weaker than the FBOE, with some exceptions (e.g., Sadr-Bazzaz & Vasey, 2025). Part of this SBOE may be spurious, due to a trivial sampling effect: in a population with an even sex-ratio, sampling individuals with more older brothers also means sampling individuals with correlatively more older sisters. This spurious SBOE has been suggested several times (e.g., Blanchard, 1997; Blanchard & Lippa, 2007), and has been formally supported by Raymond et al. (2023). However, recent evidence suggests that a real (non-spurious) SBOE also contributes to male androphilia: the number of older sisters, independent of the number of older brothers, influences the likelihood of androphilia in later-born males. This has been demonstrated by controlling for the FBOE, either experimentally by considering only specific birth ranks (Blanchard & Lippa, 2021; Khovanova, 2020), or statistically by considering large national-wide samples (Ablaza et al., 2022). This real SBOE is presently not predicted by the MIH, although it has been claimed that this SBOE is still spurious, thus still compatible with the MIH (Blanchard, 2022). The verbal argument for this claim is based on the presence of miscarriage: if some embryos are miscarried, the correlation between older brothers and older sisters is different whether all older brothers are considered (miscarried or not), or only those not miscarried. In other words, the verbal argument suggests that there is a remaining spurious SBOE after controlling for the number of older brothers that were born, due to the number of older brothers that were not born. Whether the SBOE observed is real or spurious thus remains unclear.

Second, the presence of male androphilia in male only-children, firstborn, or with only older sisters is not directly predicted by the MIH. These three categories of males do not have older brothers; their mother could thus not have developed an immune response against specific male antigens. However, there may be a natural variability in the initial maternal immune response in a population, or in alleles linked to same-sex attraction (as expected from the observed heritability for same-sex attraction, see above), so that firstborn androphilic males could emerge at a low rate. Alternatively, Bogaert & Skorska (2011) suggested that miscarried male embryos could contribute to the maternal immune response as the firstborn male is thus not necessarily the first male embryo present in the maternal womb. The relatively high estimate of miscarriage estimated in human populations (ca. 10-30%, Andersen et al., 2000; Buss et al., 2006; Wilcox et al., 1988), is consistent with this hypothesis, reinforced by the fact that miscarried male embryos are more likely to lead to immunization (Bianchi et al., 2001). However, it remains to be seen if the miscarriage rate required to generate the observed frequency of same-sex orientation in men without older brothers is quantitatively compatible with actual data on miscarriage rate.

Third, it has been observed that there is a higher prevalence of male same-sex orientation in only-children than in firstborn of larger sibships (Blanchard & Lippa, 2021), a result also found in a large sample from the Netherlands (Ablaza et al., 2022). This could be explained if there is a heterogeneity of maternal conditions: mothers who produce an androphilic son at their first delivery could belong to a biologically distinct subpopulation of mothers with a decreased probability of carrying subsequent foetuses to term (Blanchard, 2012). To explain this observation in the framework of the MIH, a different type of immune response has been hypothesized in these women, specifically affecting foetuses in first pregnancies and reducing subsequent fertility (Blanchard & Lippa, 2021; Skorska et al., 2017). However, birth weights of firstborn androphilic and gynephilic men (including only-children) are not different (Blanchard & Ellis, 2001, but see Skorska et al., 2017), a result which indirectly suggests nothing unusual about the gestation of firstborn androphilic men (Blanchard, 2012). Thus, whether an additional type of immune response, thus an extended MIH, is required, or another type of explanation, remains unclear.

Taken together, these observations and theoretical considerations highlight that the MIH provides a plausible biological mechanism for the FBOE, but its inability to fully account for the other birth order effects suggests that the hypothesis may need refinement, or that additional mechanisms are at play. To systematically evaluate these possibilities, a quantitative approach is required. By modelling the MIH and its predictions, we can explicitly test whether the observed patterns in real-world data align with its theoretical framework. This allows us to assess the necessity of extending the MIH or incorporating alternative explanations.

In this study, we addressed the following key questions. First, we developed a formal model of the MIH to: a) derive explicit functions for both the FBOE and SBOE, b) quantitatively evaluate the relative contributions of spurious and non-spurious SBOE by manipulating the correlation between the number of older brothers and older sisters, and extend this approach to other sib categories (i.e. younger brothers or sisters) as a control, c) assess the impact of miscarriage on the SBOE and on the prevalence of same-sex orientation in firstborn males, and d) compare the prevalence of male same-sex orientation between only-children and firstborns from larger sibships. This modelling approach, based on simulated data, allowed us to clarify the specific predictions generated by the MIH, and quantitatively estimate the possible importance of miscarriage for the various birth order effects. Second, using individual-level datasets from real populations, we empirically tested whether birth order effects beyond the FBOE, such as spurious or non-spurious SBOE, are consistent with the MIH when accounting for miscarriage.

## Material and methods

### Simulating random population samples

In order to evaluate the properties of family data under the MIH hypothesis, population samples of male individuals were generated, and the following sampling process was applied for each focal individual. A birth rank was drawn from the Poisson birth order distribution R(λ), where the probability of sampling an individual with birth order *j ≥* 1 is

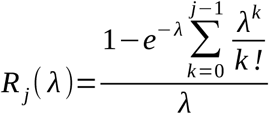

where λ is the parameter of a Poisson distribution (for details, see Raymond et al., 2023). Alternatively, when the number of younger siblings was also considered, the total number of siblings was drawn from a displaced Poisson distribution of parameter *r* = −1, with mean λ + 1 and variance λ (Staff, 1967), see Appendix 2 of Raymond et al. (2023) for details. The rank of the focal individual was randomly assigned within the sibship. When the rank of the focal individual exceeded 1, the sex of each sibling was randomly assigned under a 1.06 M/F sex ratio constraint (D’Alfonso et al., 2023) (a 1.0 sex ratio also produced qualitatively equivalent results), and the birth order was recorded. For the mother of the sibship, an initial level of immune response (*imr*) was attributed, drawn from a Gamma distribution of parameters mean = *imr_mean*, and sd = *imr_sd*. Each male birth increased *imr* by a fixed quantity (*b*): each increase of *imr* thus corresponds to an increase of the maternal immune response against male antigens. A female birth did not affect *imr* (Fig. 1). Sexual orientation for each male was determined by a logistic function of *imr:* 1/(1+*e*^−^*^slope^*^⋅^*^imr^*), with *slope* > 0 (this is an arbitrary choice among other increasing and saturating functions). Thus, as *imr* increased, the probability of a same-sex orientation increased. This process was run until the sample reached 300 gynephilic and 300 androphilic men (thus matching most empirical data sets where balanced numbers of gynephilic and androphilic male subjects are sampled from the populations), before performing the various tests and storing the corresponding estimates and *P*-values. This was replicated at least 100 times for a given set of parameters. Unless otherwise indicated, the following parameter values were considered throughout. The initial amount of immune response refers to a relative quantity and was set to an arbitrary value of *imr_mean = −5* and *imr_sd = 0.2.* The increase of *imr* for each male birth was *b* =1, and the slope of the logistic curve determining the probability of a same-sex orientation according to the amount of *imr* was set to *slope* = 1. Thus, the frequency of same-sex orientation for a firstborn male was ∼1.84 % (SEM = 1.1×10^-2^) using these standard parameters values. The mean fecundity was set at *λ* = *5,* in order to study FBOE and SBOE over a large range of birth ranks.

**Figure 1.**
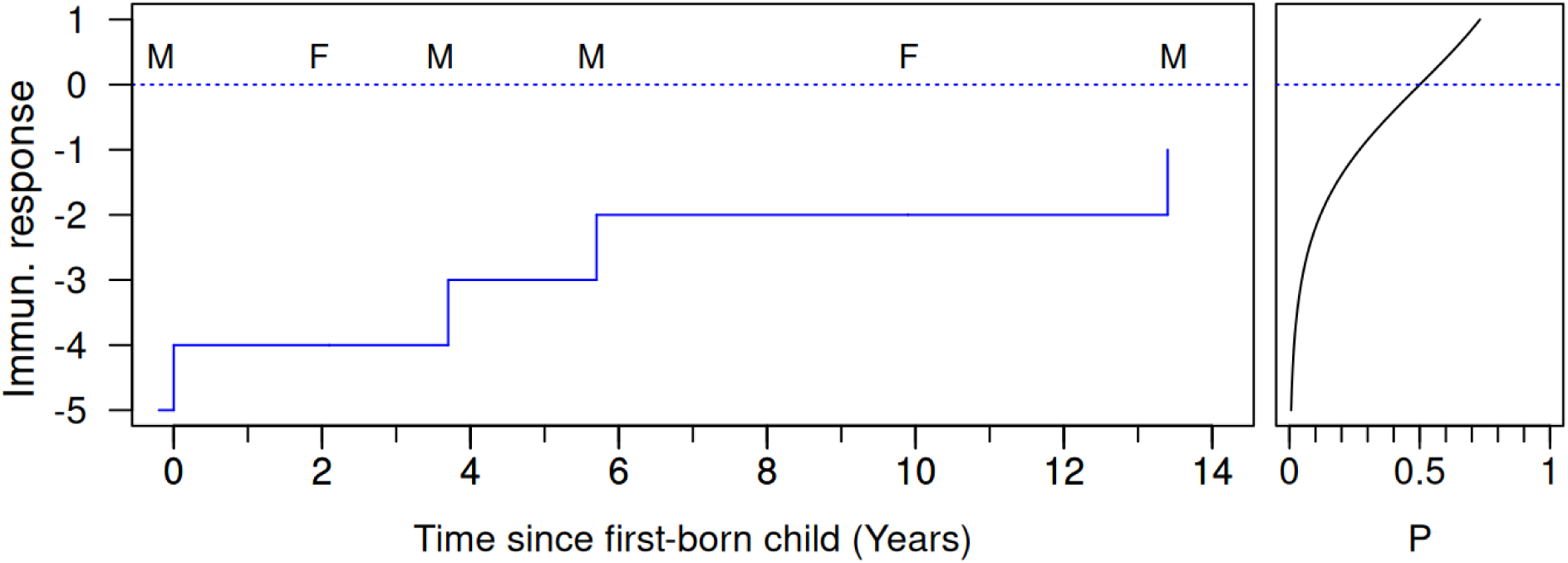
Evolution of the maternal immunological response during successive pregnancies of a mother, under MIH. Initial immune response is set at an arbitrary value (−5) before the first pregnancy. The amount of immune response increase for each male child, and is associated to the probability of a same-sex orientation (P), according to a logistic curve (right panel). The sex of each successive child (M or F) is indicated. Parameter values: increase of the *Immune response* for each son is *b* = 1. The logistic curve of sexual orientation assignation has *slope* = 1 and is centred on *Immune response* = 0 (dotted line).

In order to generate a population sample without a correlation between the number of older brothers (*ob*) and older sisters (*os*), for each individual, a male birth rank (*r_m_*) and a female birth rank (*r_f_*) were independently drawn from the rank sampling distribution R(λ/2) (see Eq A1b.1 of Raymond et al., 2023). The resulting number of older brothers (*r_m_ - 1*) and older sisters (*r_f_ - 1*) were added, and their birth order was randomly assigned before calculating *imr* levels, and sexual orientation, as above.

In models that incorporate the effect of miscarriage, siblings were assumed to be subject to a miscarriage with probability *fm* (ranging from 0 to 0.7) affecting both male and female embryos (*fm,* or frequency of miscarriage, represents the probability of miscarriage for each pregnancy). Each miscarried older sib increased the level of *imr* by *b*(1−*α*) for a boy (no change for miscarried girls), where *α* ∈[0, 1].

### Statistical analysis of simulated data

The birth order effect, i.e., the increase in the probability *p* of displaying a same-sex orientation with the number of older siblings was modelled as *p* = *f(X)* where *X* is the number of older brothers (for modelling FBOE), or the number of older sisters (for modelling SBOE). The function *f* was inferred from modelling (a logistic function for FBOE, see below) or, when such inference was not possible (for SBOE), various continuous forms of the function *f* were considered, notably logistic, logistic with a polynomial effect, saturating exponential, or geometric (Table S1). These functions describing the SBOE effect were compared by fitting models to the simulated data using MCMC, then comparing the various SBOE functions using WAIC (Watanabe-Akaike Information Criterion), a generalized version of AIC onto singular statistical models, see Gelman et al. (2014) and Watanabe (de Valpine et al., 2020; 2013): the mean of ten independent chains was used, each with a length of 50,000 samples and a burn-in phase of 20,000. To avoid the effects of small sample sizes for the number of older sisters, we restricted the data to categories of birth rank including at least 50 individuals.

The presence of a possible spurious SBOE was assessed using two different approaches for controlling for a FBOE. First, a generalized linear model with the sexual orientation as the response variable, the number of older sisters (*os,* quantitative) as the variable of interest, and the number of older brothers (*ob,* quantitative), and the number of sibs (*sib*, quantitative) as control variables, as in (Blanchard et al. 2025). Second, by considering only individuals with a specific male birth rank (here male birth rank = 1, i.e. *ob* = 0) and performing a generalized linear model with the sexual orientation as a response variable, and *os* and *sib* as independent variables. In both cases, the significance of the variable of interest (*os*) was calculated by removing it and comparing the resulting variation in deviance using the χ2 test, as done by the function Anova from the R package *car (Fox & Weisberg, 2019)*. The frequency of significant SBOE was calculated on the results of at least 400 independent samples. Significant *P*-values indicative of a SBOE corresponded to cases where more older sisters were associated with a same-sex orientation, relatively to heterosexual men (i.e., a positive slope estimate of the variable of interest), so the expected false discovery rate was 2.5% (one-sided test).

### Estimating the miscarriage frequency from simulated data

To estimate the miscarriage frequency from the observed SBOE, a population sample was generated with a specific frequency of miscarriage *fm*, a mean observed fertility *λ*_obs_ = 3 (thus a real mean fecundity of *λ*_obs_/(1-*fm*)), and composed of the same number of androphilic or gynephilic men, with the total size (androphilic plus gynephilic) being *N*_obs_ = 2,400. The effect of the number of older sisters (*os*) on sexual orientation was tested with a logistic regression, with the number of older brothers and the number of sibs as independent variables, and the estimated slope associated with *os* (*slope_obs*) was used to estimate *fm*. This was done using *fm_estim(freq_mis)*, a function with a parametric value of frequency of miscarriage *freq_mis*, generating a population sample with the same observed fertility as in the sample, the same proportion of androphilic or gynephilic men, with a total size of *n*.*N*_obs_ (with *n* = 20, *n* being an amplification parameter reducing the variance of the estimates; lower values of *n*, including *n* = 1, provided quantitatively similar mean estimates), and returning the absolute difference between the calculated slope of SBOE and *slope_obs*. The value of *freq_mis* generating the minimum of the function was calculated using *optimize()* from the stats R package. This process was replicated at least 200 times for each *fm* value, for *fm* value from 0.1 to 0.7.

### Empirical individual family data

Sampling in French Polynesia was conducted in January and October 2023, and May 2024, using a snowball sampling approach. Participants were recruited in public areas (e.g., markets, malls, parks) and private locations (e.g., hotels, shops) on the islands of Tahiti, Moorea, and Bora-Bora. Prior to participation, each individual received a document outlining the study’s purpose and the contact details of the principal investigator (M.R.). This document explicitly states that personal data will only be used for research purposes and that only global results –not individual data– will be published. Written informed consent was obtained from all participants. This project was approved by the ethical committee of the University of Montpellier (n°UM 2023-008bis), and protocols used to recruit individuals and to collect data were approved by the French National Committee of Information and Liberty (CNIL) through the CNRS (approval #1226659). Each participant was interviewed confidentially and anonymously at a convenient location near the point of contact. They were asked to report their sex assigned at birth (male or female), gender identity (male, female, or third gender such as Rae rae or Mahu), date of birth, self-declared sexual orientation (attraction to males, females, both, or other), and the island or country of birth of their four grandparents. Individuals below 18 years of age (legal age of majority in France) were not considered. To reduce cultural heterogeneity, individuals with three or more grandparent(s) born outside Polynesia were not further considered. Individuals from multiple births, or with brothers of sisters from multiple births, were also not considered. In each sibship, full-sibs or half-sibs with the same mother as the focal individual were kept. Usable data were obtained from *N* = 380 men of mean age (M +/−SEM) = 34.8 +/− 0.60, including *N* = 149 androphilic (mean age = 31.6 +/−0.78, mean date of birth = 1992.2), and *N* = 231 gynephilic (mean age = 37.0 +/− 0.82, mean date of birth = 1986.8). Additional datasets were selected based on two explicit criteria: (1) the sample included both male heterosexual and homosexual individuals, and (2) sibling data were available, specifically the number of older and younger brothers and sisters. All accessible datasets meeting these criteria were included: 1) the Indonesian sample used in Nila et al. (2019), and fully described in Nila et al. (2018), and restricted to men with either a homosexual or heterosexual orientation (thus removing the “Bisexuals” category) 2) the French sample described in Raymond et al. (2023), restricted to men with either a homosexual or heterosexual orientation (thus removing the “Bisexuals” category), 3) the Canadian sample from Blanchard and Bogaert (1996), 4) the Samoa sample from Semenyna et al. (2023), 5) The Greek sample described by Apostolou (2020a), restricted to men with either a homosexual or heterosexual orientation (thus removing “Bisexuals” and “Heterosexual with same-sex attractions”), 6) the Czech sample from Fořt et al. (2024), restricted to men, and used without imputing missing values, and 7) the Iranian sample from Sadr-Bazzaz and Vasey (2025), restricted to individuals whose sex assigned at birth was male (thus including transgender males), and whose sexual orientation was either androphilic or gynephilic (thus removing the ambiphilic category). The basic sibship data for these eight datasets are detailed in Table S3.

### Statistical analysis of empirical individual family data

The presence of an FBOE and SBOE in the population samples was assessed using an ABZ regression (Blanchard et al., 2025; Zdaniuk et al., 2025), a generalized linear model with the sexual orientation as the response variables, the number of older sisters (*os,* quantitative), the number of older brothers (*ob,* quantitative), and the total number of sibs (*sib*, quantitative) as independent variables. When the slope of *os* (*slope_os*) was positive (corresponding to a SBOE), the amount of miscarriage (*fm*) required to generate it was estimated using the following procedure, for each population sample *i*. First, the demographic parameters potentially influencing the correlation between *ob* and *os*, i.e. the mean and variance of the observed fertility, were calculated: the mean fertility *λ_i_*was estimated from the heterosexual sample only (as the mean fertility observed in samples of same-sex males is slightly inflated when there is a FBOE due to their higher mean birth rank), using the mean number of brothers and sisters, see Raymond et al. (2023) for details; the variance in fertility was modelled as *λ_i_/ν_i_*, with *ν_i_* measuring over- or under-dispersion relatively to a Poisson distribution (*ν_i_*< 1, or *ν_i_* > 1, respectively), and *ν_i_*was estimated using the COMPoissonReg R package (Sellers & Shmueli, 2010). Second, *fm_i_* was estimated using *fm_estim2(freq_mis)*, a function with a parametric value of frequency of miscarriage *freq_mis*, generating a population sample with the same parameters *λ_i_*, and *ν_i_*, the same proportion of androphilic and gynephilic men as observed in population sample *i*, with a total size of *n*.*N*_obs_, (with *n* = 50, *n* being an amplification parameter reducing the variance of the estimates; lower values of *n*, including *n* = 1, provided quantitatively similar mean estimates, see Fig. S3) and calculating the slope of *os* from an ABZ regression, and returning the absolute difference between this calculated slope and *slope_os_i_*. The value of *freq_mis* generating the minimum of the function was calculated using *optimize()* from the stats R package. This process was replicated at least 200 times, and the mean of the replicates was used as the estimated *fm_i_*.

The presence of a younger sib effect was assessed using a variation of the ABZ regression, with the sexual orientation as the response variables, the number of younger brothers (*yb*, quantitative), the number of older brothers (*ob*, quantitative), and the number of sibs (*sib*, quantitative) as independent variables. Another regression was similarly performed by replacing the number of younger brothers by the number of younger sisters (*ys*, quantitative),.

All analyses were run in R version 4.5.2 (R Core Team, 2025)

## Results

### Modelling the FBOE generated by MIH

Let’s consider an initial immunological maternal response level of *x_0_*. We assume that, under MIH, only sons increase the level of *imr* (*i.e.,* increase the specific immune response), with each son increasing this *imr* by a quantity *b*, with *b > 0* (Fig. 1). The level of *imr* is *x_1_= x_0_ + b* for a firstborn male. Successive brothers have an additive effect on the amount of *imr*: *x_n_ = x_0_ + b.n*, or *x_n_* = *x_0_ +b + b.ob*, where *n* is the male birth rank and *ob* the number of older brothers (*ob = n - 1*). The probability of a same-sex orientation as a function of the number of older brothers is given by *p_n_*=*f* (*x_n_*), where *f* is a logistic function with a positive slope (so that *p_n_* increases as x_n_ increases, see Fig. 1).

Simulations were conducted by varying two key parameters: *b* (the additional immune response triggered by a male gestation) and the slope of the logistic curve (which determines the probability of a same-sex orientation based on the immune response level). This generated a FBOE described by a logistic regression: when the effect on the immune response of each older brother was relatively large (e.g., *b* > 2), the proportion of androphilic men *p_n_* reached a plateau at *p_n_*≈1 for three or four older brothers. For relatively smaller effects of each older brother (e.g., *b* < 2), *p_n_*displayed a continuous and sigmoid increase over the common range of 0-4 older brothers (Fig. 2A). The same effect was observed when *slope* varied (i.e., the slope of the logistic function determining sexual orientation from the level of the immunological response), with higher values decreasing the proportion of androphilia in individuals with a low number of older brothers and increasing those with higher numbers of older brothers (Fig. 2B).

**Figure 2.**
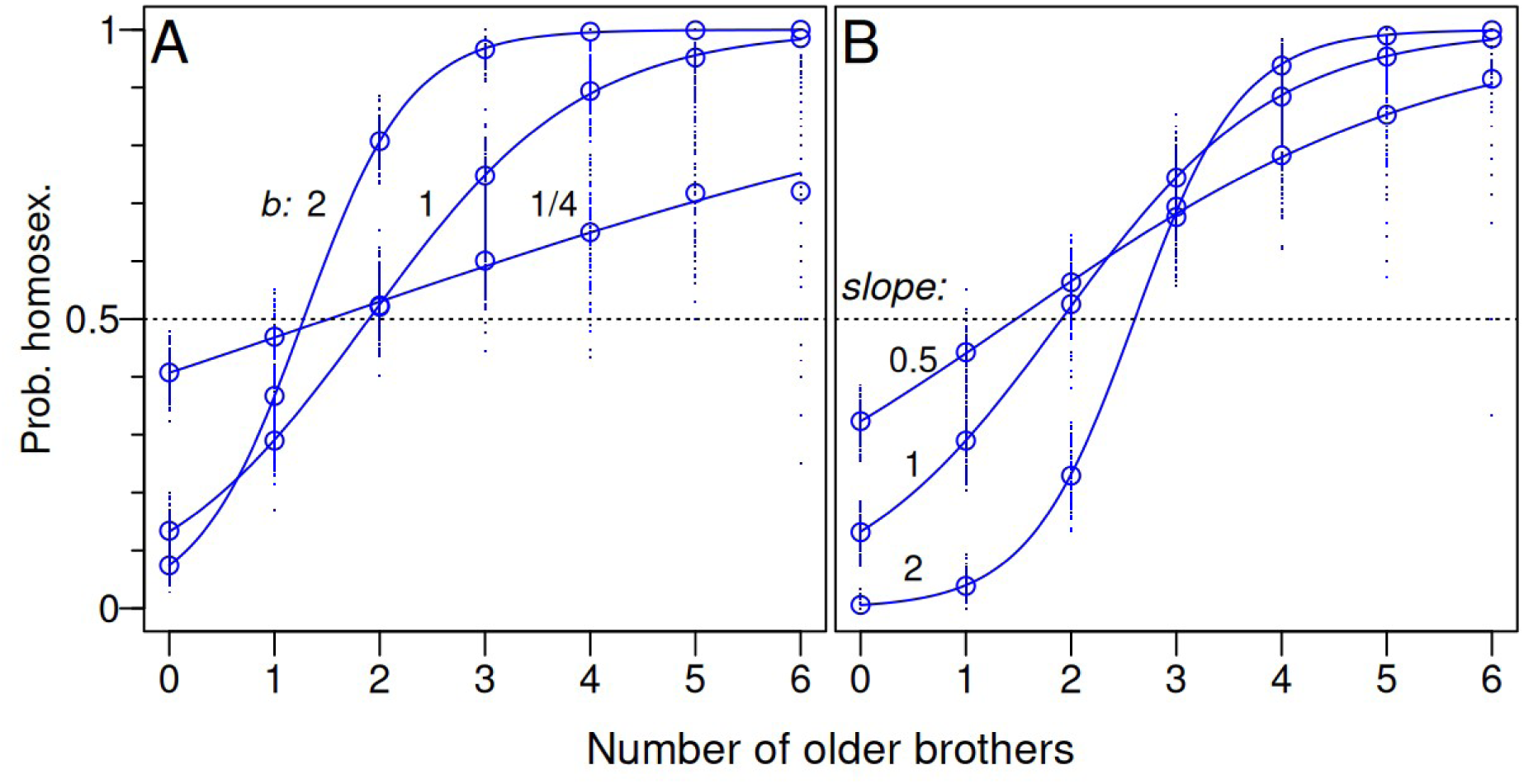
FBOE generated by the MIH. Simulating random population samples of gynephilic and androphilic men with a variable value of *b* (A), or *slope* (B), and computing the proportion of androphilic men for each number of older brothers. The increase of *imr* for each son is *b* = 1/4, 1, or 2, with *slope* = 1 (A), and the slope takes the values 0.5, 1, or 2, with *b* = 1 (B). The mean of 100 replicates, for each number of older brother, is depicted as a circle. Lines are the expected curve from a logistic regression. Absence of FBOE is indicated by a dotted line, corresponding to *b* = 0 (A), or *slope* = 0 (B), at the level (0.5) corresponding to the sample frequency of androphilic men.

### Spurious SBOE

In addition to a FBOE, a SBOE was apparent when analysing simulated data under the MIH (Fig 3). The theoretical shape of this SBOE was not readily identifiable; various functions (Table S1) describing the variation of probability of a same-sex orientation according to the number of older sisters were thus compared using WAIC. This SBOE was best described by a saturating exponential with 2 or 3 parameters, particularly when the shape of the curve can be estimated over a large range of number of older sisters (e.g., up to 6 older sisters). When a more restricted range of number of older sisters are available (for example when having a limited sample size from a population with a low mean fertility), a larger number of possible functions could describe this SBOE, including a logistic function (Table S2).

**Figure 3.**
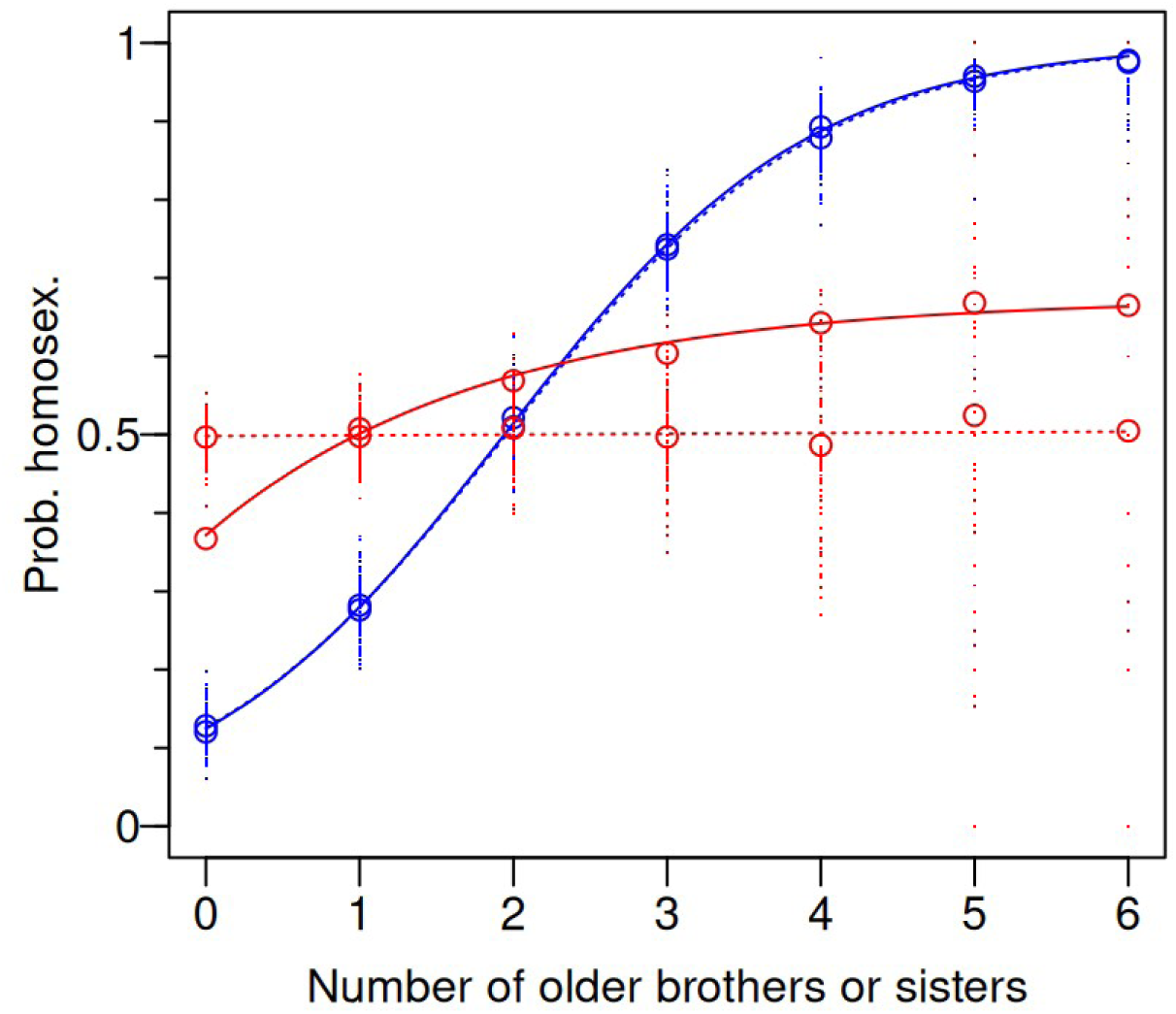
Effect of the correlation between the number of older brothers and older sisters on FBOE and SBOE. The curves depict the predicted proportion of androphilic men according to the number of older brother (FBOE, in blue), or older sisters (SBOE, in red). Data contained the same number of androphilic or gynephilic men (thus a frequency of androphilic men of 0.5), and a total size being N ≈ 600, from a population with mean fertility l = 5. They were generated with (plain line) or without (dotted line) a correlation between older brothers and older sisters. The coloured circles represent the mean of up to 100 replicates, with the same colour code as above. For clarity, these replicates are depicted as coloured dots only for the data generated with uncorrelated *ob* and *os*.

This SBOE is necessarily spurious as, under our simulation of the MIH, older sisters have no influence on the sexual orientation of their younger brothers since they have no direct effect on the maternal immune response. When data were generated without a correlation between the number of older brother and older sisters (by generating independently the number of older brothers and older sisters), the FBOE was not significantly affected: the slope decreased from 1.004 (linear unit) to 0.990, and this change was not significant (*X^2^*= 1.4 10^-3^, *df* = 1, *P* = 0.242). However, the SBOE became not significantly different from a line of slope = 0 (*X^2^* = 0.38, *df* = 1, *P* = 0.54), suggesting that this spurious SBOE is the result of the correlation between the number of older brothers and older sisters (Fig. 3). Indeed, when this correlation is taken into account in a logistic regression, i.e., when *ob* is introduced as a control variable, the proportion of significant SBOE was not different from the false rejection rate of 2.5%, for all sample sizes (Binomial test, *P* = 0.074, 0.074, and 0.333, for sample sizes N = 600, 1,200, and 2,400, respectively).

However, when miscarriage was present and increased in frequency, the rate of false rejection increased, and became significantly above the false rejection rate of 2.5 % (Binomial test, *P* < 10^-4^) from a miscarriage rate of 5%, for all sample sizes (Fig. 4A). This increase of false rejection rate was more pronounced for higher sample size values. To better understand this result, three additional analyses were performed. 1) The logistic regression was performed on a restricted data containing only the firstborn among the males (i.e., *ob* = 0, and *os* ≥ 0), and with only the number of sibs as a control variable: the results were similar but less pronounced (Fig. 4B). 2) Samples were generated without the possibility for female embryos to be miscarried, thus there was no correlation between the number of miscarried male and female embryos: the results were similar but more pronounced (Fig. S4). 3) Samples were generated without a correlation between the number of older brothers and the number of older sisters, and in presence of miscarriage: the rate of false rejection was not different from 2.5%, for all sample sizes (across all miscarriage frequencies, Fisher’s method, *P* = 0.925, 0.811, or 0.250, for N = 600, 1,200, or 2,400, respectively). 4) When the number of older brothers was replaced, in the regression, by the initial number of older brothers (before miscarriage), the proportion of significant SBOE was not different from the false rejection rate of 2.5%, for all sample sizes (across all miscarriage frequencies, Fisher’s method, *P* = 0.925, 0.852, or 0.946, for N = 600, 1,200, or 2,400, respectively).

**Figure 4.**
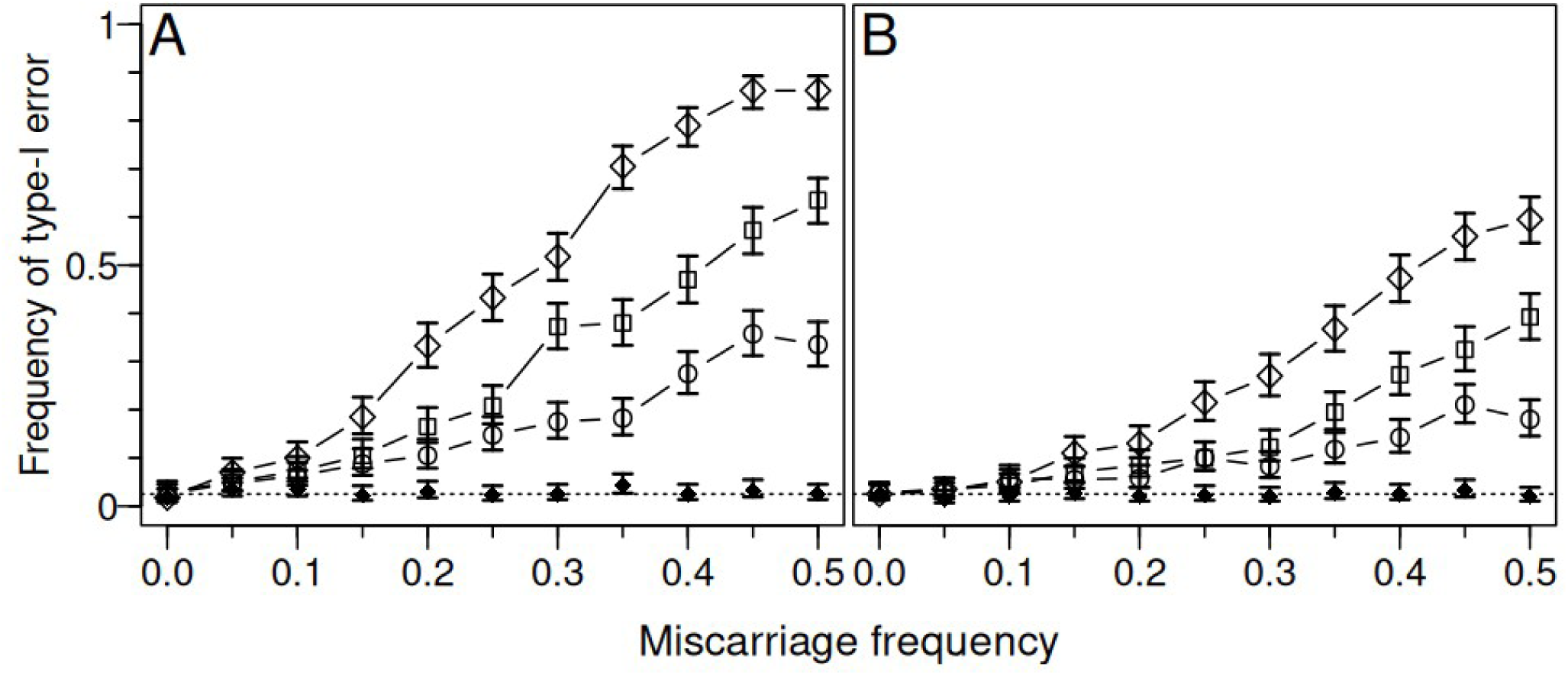
Effect of miscarriage frequency on type-I error for detecting an SBOE. Data were generated under the MIH for various values of miscarriage affecting both male and female embryos, and SBOE was detected using a logistic regression, controlling for the number of older brothers statistically (A), or by restricting the data to firstborn males (B). Each point depicts the proportion of significant SBOE over 400 independent replicate samples, with the corresponding 95% confidence interval. Each sample is composed of the same number of androphilic or gynephilic men, with the total size being N = 600 (empty circles), N = 1,200 (empty squares), or N = 2,400 (empty rhombus). The proportion of significant SBOE when controlling for the initial number of older brother (before miscarriage), or by restricting the data to firstborn among all males (before miscarriage), is depicted by full small rhombus, in A, and B, respectively (only N = 2,400 is shown). The dotted horizontal line indicates the false rejection rate of 2.5%.

A younger brother effect was apparent when analysing simulated data under the MIH: same-sex-oriented men tend to have significantly fewer younger brothers (mean slope over 400 independent samples = −0.36, SE = 0.004; all *P* < 0.05). To better understand this result, data were generated without a correlation between the number of older and younger brothers (by generating independently the number of older and younger sibs): the proportion of significant younger brother effect was not different from the false rejection rate of 2.5% (Binomial test, *P* = 0.333, for sample sizes N = 600), suggesting that this spurious younger brother effect is the result of the correlation between the number of older and younger brothers. Indeed, when this correlation is taken into account in a logistic regression, i.e., when *ob* is introduced as a control variable, the proportion of significant younger brother effect was not different from the false rejection rate of 2.5% (Binomial test, *P* = 0.747, for sample sizes N = 600). The correlation between older and younger brothers was found negative (mean Kendall correlation over 400 independent samples = −0.295, SE = 0.001). Similar results were found when the number of younger brothers was replaced by the number of younger sisters (not shown). When miscarriage was present and increased in frequency, the rate of false rejection increased, despite the presence of *ob* as a control variable, and became significantly above the false rejection rate of 2.5 % (Binomial test, *P* < 10^-2^) from a miscarriage rate of 10%, for all sample sizes. This increase of false rejection rate was more pronounced for higher sample size values (Fig. S5). When the number of older brothers was replaced, in the regression, by the initial number of older brothers (before miscarriage), the proportion of significant younger sib effect was not different from the false rejection rate of 2.5% (across all miscarriage frequencies, Fisher’s method, *P* = 0.706, for N = 2,400, Fig. S5).

In presence of miscarriage, the slope of the regression corresponding to SBOE was used to estimate the frequency of miscarriage *fm*. Negative slopes were not considered. The mean estimated values of *fm* were linearly related to the true *fm* values (Fig. S1), with the slope of the regression line (slope = 0.88, SEM = 0.052) marginally not significantly different from 1 (*t* = −2.338, *P* = 0.067). When the point corresponding to *fm* = 0.1 was omitted, the slope was not significantly different from 1 (*t* = −1.632, *P* = 0.178) and was closer to 1 (slope = 0.96, SEM = 0.022), suggesting that *fm* is slightly overestimated for low values of *fm*, probably due to the impossibility to estimate *fm* when the observed SBOE slope is negative.

### Firstborn androphilic men and miscarriage

On simulated data, the percentage of firstborn male individuals displaying same-sex orientation (including only-children) was not constant when the frequency of miscarriage increased. For a given mean fecundity λ, increasing the frequency of miscarriage *fm —*thus with a decreasing observed mean fertility λ(1-*fm*)— increased the frequency of male same-sex orientation among firstborn individuals (Fig. S2, A). For the given observed mean fertility λ, increasing the frequency of miscarriage *fm —*thus with an increasing mean fecundity λ/(1-*fm*)— significantly decreased (β = −0.028, F_1,7_ = 8.95, *P* = 0.020) the frequency of male same-sex orientation among firstborn individuals (Fig. S2, B).

In the absence of miscarriage, the prevalence of male same-sex orientation in only-children and firstborn of larger sibship was not statistically different (Fisher exact test on 2×2 contingency table, *P* = 0.323). When the frequency of miscarriage increased, the prevalence of male same-sex orientation became significantly higher in only-children relatively to firstborn of larger sibship (10% miscarriage: *P* = 0.034; 20% miscarriage or above: *P* < 10^-10^). This effect was observed for a large range of lambda values, except when lambda was very low (Fig. 5).

**Figure 5.**
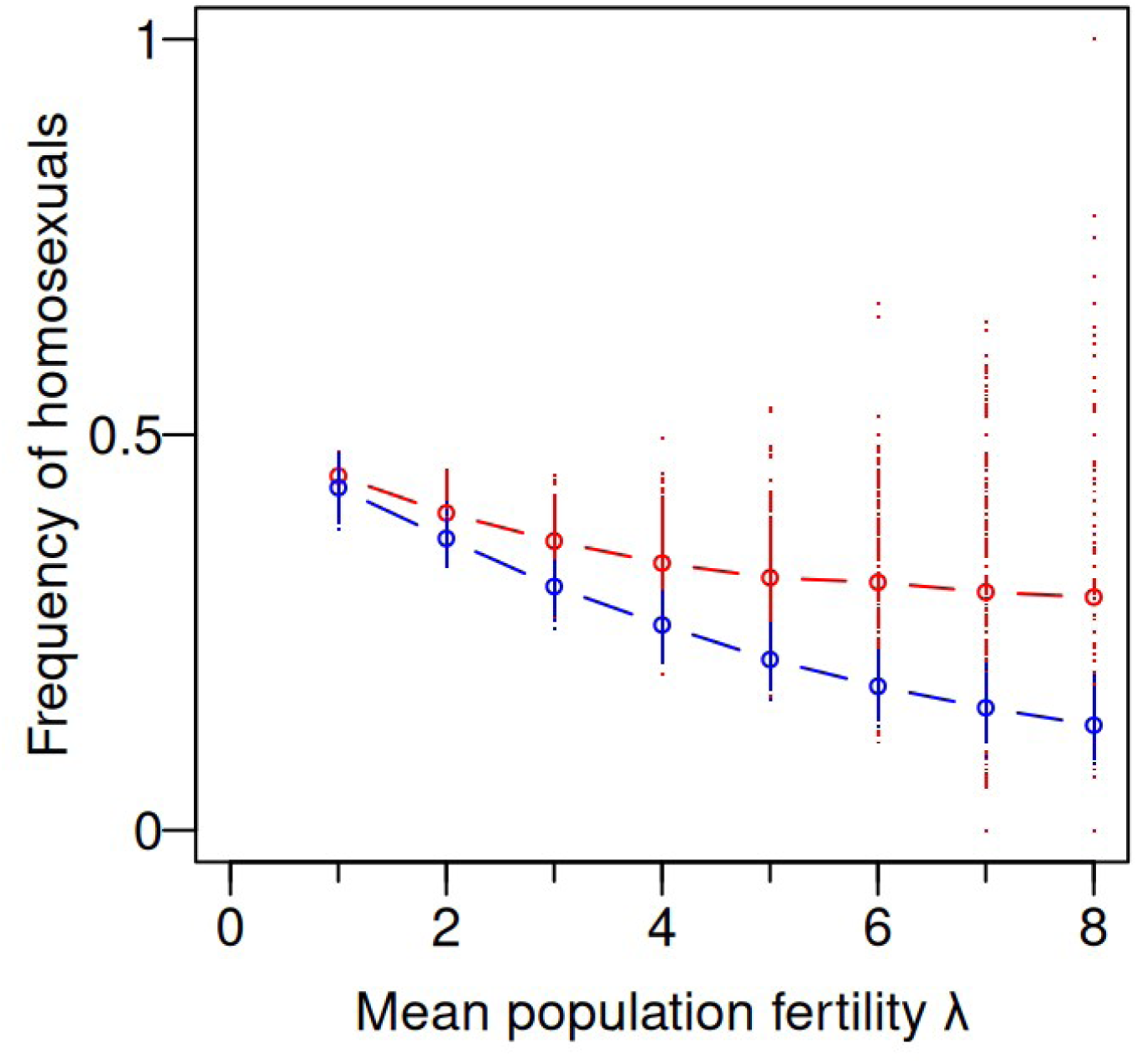
Effect of the mean population fertility on the frequency of androphilic men in only-children and firstborn with younger siblings. Data are generated using a mean fertility of *λ* = 5, and a rate of miscarriage *fm*= ½, thus with an observed fertility of *λ(1 - fm)*, and with a miscarriage cost α = 25%. Each sample is composed of the same number of androphilic or gynephilic men, with the total size being N = 2,400. The frequency of androphilic men in only-children and in firstborn with younger siblings are depicted in red and blue dots, respectively. The coloured circles, with the same colour code, represent the mean of at least 200 replicates for each value of lambda.

### Empirical population data

For the various empirical population datasets, the correlation between *ob* and *os* (τ_ob-os_) varied from 0.007 to 0.385, with a mean of 0.199 (SEM = 0.050), significantly different from zero (Wilcoxon signed rank exact test, *P* = 0.008). All sample values were significantly higher than 0, except the French dataset (Table S3). The variability of τ_ob-os_ values was well explained by the mean fecundity *λ* and the under/over-dispersion *ν* parameter: a linear regression model using *λ* and *ν* to predict *τ_ob os_* explained 95.7% of the deviance, and the higher the mean fertility (higher *λ*), and the more over-dispersed the fertility distribution (lower *ν*), the higher the value of τ_ob-os_ (mean fertility *λ*: *β* = 0.059, SEM = 0.013, *X^2^* = 21.7, *df* = 1, *P* = 3.2×10^-6^; under/over-dispersion *ν* parameter: *β =* −0.127, SEM = 0.039, *X^2^*= 10.6, *df* = 1, *P* = 1.1×10^-3^).

Overall, the number of older sisters (*os*) had a significant effect on the sexual orientation, when the number of older brothers, and the sib size, were statistically controlled for (*X^2^*= 32.1, *df* = 1, *P* = 1.5×10^-8^), Table 1. With the full model incorporating also intercept and slope variation across populations (i.e. *pop* and *pop:os*), *os* was significant (*X^2^* = 19.5, *df* = 1, *P* < 1.02×10^-5^), as well as the intercept *pop* (*X^2^*= 309.9, *df* = 7, *P* < 10^-10^) reflecting different frequencies of androphilic men across the dataset, and the interaction *os:pop* (*X^2^* = 14.1, *df* = 7, *P* = 0.049), marginally suggesting distinct slopes for *os* across the datasets. To remove directly the correlation between older brothers and older sisters, data were restricted to individuals with a fixed number of older brothers, the largest category being individuals without older brothers (*ob* = 0). The number of older sisters still had a significant effect on the sexual orientation (*X^2^* = 11.8, *df* = 1, *P* = 5.9×10^-4^). The slope of *os* was statistically not different across the datasets (interaction *os:data* : *X^2^* = 5.75, *df* = 7, *P* = 0.569).

**Table 1.** Slopes of the binomial ABZ regression from population data. The slope (β) of *ob* (for FBOE) and *os* (for SBOE) are given, with their corresponding SEM and *P*-value, for each sample. For positive slope of SBOE, the frequency *fm* of miscarriage required to generate the SBOE slope is given (distribution of *fm* in Fig. 6). ‘All’ refers to values generated when all population samples were pooled. Significant (*P* < 0.05) *P*-values are in bold.

| Sample | FBOE |  |  | SBOE |  |  | Miscarriage |  |
| --- | --- | --- | --- | --- | --- | --- | --- | --- |
| | $\beta$ | (SEM) | P-value | $\beta$ | (SEM) | P-value | *fm* | (SEM) |
| Canada | 0.324 | (0.103) | **0.001** | 0.134 | (0.098) | 0.172 | 0.365 | (0.001) |
| Czech | 0.405 | (0.093) | **<10<sup>-4</sup>** | 0.195 | (0.091) | **0.032** | 0.569 | (0.001) |
| France | 0.390 | (0.157) | **0.012** | 0.163 | (0.146) | 0.261 | 0.456 | (0.003) |
| Greece | 0.364 | (0.141) | **0.010** | 0.213 | (0.149) | 0.156 | 0.577 | (0.001) |
| Indonesia | 0.164 | (0.174) | 0.336 | 0.015 | (0.158) | 0.926 | - | - |
| Iran | 0.303 | (0.125) | **0.013** | 0.263 | (0.122) | **0.030** | 0.566 | (0.001) |
| Polynesia | 0.200 | (0.101) | **0.047** | -0.003 | (0.107) | 0.975 | - | - |
| Samoa | 0.177 | (0.054) | **0.001** | 0.182 | (0.057) | **0.001** | 0.401 | (0.001) |
| All | 0.263 | (0.033) | **<10<sup>-10</sup>** | 0.189 | (0.034) | **<10<sup>-7</sup>** | 0.480 | (0.001) |

Seven population samples displayed a positive slope associated to *os*, indicating either a spurious SBOE resulting from miscarriage, or (non-exclusively) a true SBOE resulting from a distinct mechanism than MIH. Assuming the absence of the latter, the miscarriage frequency *fm* required to explain the apparent SBOE was estimated, except when slopes were low (Indonesia, slope = 0.015), in order to avoid overestimation for *fm* (Fig. S1). The resulting *fm* estimates (Table 1) were between 0.365 (SEM = 0.001) for the Canada sample, and 0.577 (SEM = 0.001) for the Greece sample, displaying unimodal distributions (Fig. 6), with a decreased variance when the amplification parameter *n* increased (Fig. S3). Over all samples, *fm* was estimated at 0.480 (SEM= 0.001).

**Figure 6.**
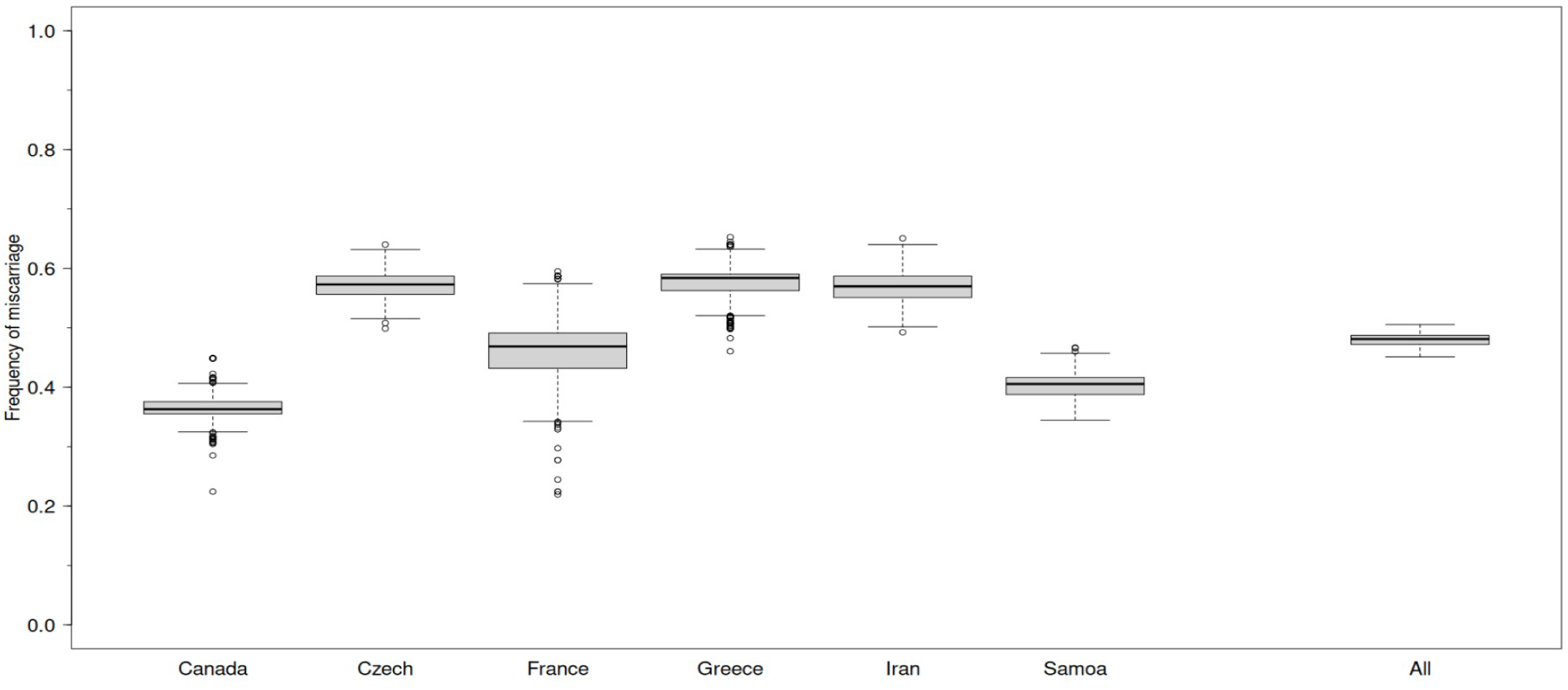
Estimating the frequency of miscarriage (*fm*) from the slope of SBOE, for each population sample. The effect of the number of older sisters on sexual orientation was tested with a logistic ABZ regression, with the number of older brothers and the number of sibs as control variables, and the corresponding slope was used to estimate *fm* by minimizing the function *fm_estim2()*, for each population sample, or for all population samples pooled (“All”). Amplification parameter: *n* = 50 (see Figure S3 for other values). The distribution of at least 200 replicates is shown. See Table 1 for mean values.

In all population samples, the frequency of same-sex orientation was higher in only-children than in firstborn with younger siblings, except the Indonesian sample displaying a low number of only-children (N = 4). This difference was significant (Fisher’s test on 2×2 contingency table, one-tailed, *P* < 0.05) for three population samples, and marginally non-significant (0.05 < *P* < 0.10) for two population samples (Table S4). Across all population samples, the frequency of same-sex orientation was higher in only-children (*f_H_* = 0.459) than in firstborn with younger siblings (*f_H_* = 0.363), the difference being significant (combined *P*-values using Fisher’s method: *P* = 0.002). This difference suggests an impact of miscarriage.

The mean correlation between *ob* and younger brothers, or younger sisters, was −0.169 (SEM = 0.022), or −0.168 (SEM = 0.015), respectively. Overall, the number of younger brothers had a significant effect on the sexual orientation, when the number of older brothers, and the sib size, were statistically controlled for (*X^2^* = 7.45, *df* = 1, *P* = 0.006): more younger brothers was associated with less same-sex oriented men (Table S5). Similarly, the number of younger sisters had a significant effect on the sexual orientation (*X^2^* =15.8, *df* = 1, *P* = 7.2×10^-5^): more younger sisters was associated with less same-sex oriented men (Table S5).

## Discussion

The aim was to evaluate if the various birth order effects other than the FBOE can be accounted for by the MIH, or if additional mechanisms are to be considered. To that aim, we modelled the MIH and derived explicit functions for the FBOE and SBOE to identify their predicted shapes, and we showed that the SBOE is non-significant when data are generated without the correlation between older brothers and older sisters, or when this correlation is controlled for in statistical analyses. However, in the presence of miscarriage, the statistical control of this correlation is insufficient and a significant spurious SBOE is apparent in a regression, with or without restricting the data to a specific birth order. Additionally, and counter-intuitively, miscarriage increases the prevalence of male same-sex orientation in only-children compared to firstborn of larger sibships. Analysing empirical data from different populations data showed the presence of a significant SBOE, and an overall higher frequency of same-sex orientation in only-children relatively to firstborn of larger sibships, compatible with the effect of miscarriage. However, the estimated level of miscarriage required to explain the SBOE observed in real population data seems relatively high compared to the published rates of miscarriage, suggesting that some parameters were not taken into account, or that additional hypotheses, beyond the MIH, are required to explain these observations.

The classical Maternal Immune Hypothesis was initially proposed to account for the observation that the presence of older brothers increases the odds of same-sex orientation, while the presence of older sisters or younger sibs does not (Blanchard & Bogaert, 1996). Under this hypothesis, older brothers have a cumulative influence on the neuro-development of their younger brothers through long-lasting effects on the immune system of their mother as it reacts to specific male antigens. This “older brothers effect” has long been suspected to generate an apparent older sister effect, due to the correlation between the number of older brothers and the number of older sisters in population data (Blanchard, 1997). It is surprising to note that, to our knowledge, this correlation has never been reported over the past 3 decades of studies addressing the FBOE or SBOE.

Here, we could verify that the number of older brothers and older sisters are positively correlated by simulating virtual population samples with fertility values similar to those of human population; we also found this correlation in most empirical human population samples analysed. In spite of this correlation, for a given number of older sibs, the numbers of older sisters and older brothers are negatively correlated (more older brothers means fewer older sisters). The variance in sibship size generates a variance in number of older sibs, with a resulting positive correlation between *ob* and *os*, this correlation increasing with the mean fertility (and thus the variance), and also with the overdispersion of the variance relatively to a Poisson distribution.

Several solutions can be employed in analysis of empirical population data to protect from the correlation between *ob* and *os* and examine the SBOE independently of the effects of older brothers. The first one is to evaluate the SBOE while controlling for the number of older brothers, thus statistically controlling for the correlation between older brothers and older sisters, as suggested by e.g., Blanchard (1997). Another option is to restrict the data to a specific male birth rank, which allows also to control for the correlation (Blanchard & Lippa, 2021). The first report of an SBOE, despite such a control, was published in 2005 (King et al., 2005). Since then, various studies from distinct populations reported also a significant SBOE while controlling for FBOE (e.g., Vasey & VanderLaan, 2007; VanderLaan & Vasey, 2011; Blanchard & Lippa, 2021; Ablaza et al., 2022; Kabátek & Blanchard, 2024; Sadr-Bazzaz & Vasey, 2025). To reconcile this apparently genuine SBOE with the MIH, it has been proposed that miscarried male foetuses, contributing to the building up of maternal antibodies and thus to FBOE, were not properly taken into account and thus that the resulting significant SBOE, controlling for the born older brothers, was still spurious.

To examine this hypothesis, we simulated data with various frequencies of miscarriage and confirmed that miscarriage can generate a spurious SBOE in a regression model when the number of older brothers is controlled for. Interestingly, this spurious SBOE vanishes if the total number of older brothers (miscarried or not) are used, instead of the live-born ones, in the regression, or if the data are generated without a correlation between the number of older brothers and older sisters. This suggests that removing randomly sibs, including older sibs, affects the correlation between older brothers and older sisters, rendering the use of the live-born older brothers as a control variable insufficient to control for the spurious SBOE. Interestingly, the correlation between miscarried male and female embryos does not participate in this effect, as the spurious SBOE is not diminished when miscarriage is restricted to only male embryos. Similarly, restricting the dataset to a specific male birth rank (and considering only live-born sibs) is also insufficient to control for the spurious SBOE. This phenomenon would apply to population samples if miscarriage were present, suggesting that when a significant SBOE is observed, it is not possible to disentangle the effect of miscarriage from a putative real SBOE, even when male birth rank is controlled for.

As a consequence, a spurious SBOE is generated when there is an FBOE, due to the positive correlation between older brothers and older sisters. Interestingly, older brothers are negatively correlated with younger brothers (or younger sisters), so a FBOE produces also a spurious younger sibling effect, with an opposite direction: same-sex-oriented men tend to have fewer younger brothers or younger sisters. Controlling for male birth rank, and thus for the correlations with older brothers, eliminates these spurious effects, unless substantial miscarriage occurs.

In simulated data, the percentage of androphilic men among the firstborn (including only-children) did not vary clearly with the miscarriage rate (Fig. S2). The theoretical sample frequency of firstborn individuals (*f _i_*=(1−*e*^−^*^λ^*)/ *λ*, see Raymond *et al*. 2023) decreases when the mean fertility λ increases, and this *f_i_*frequency increases when the miscarriage rate increases, for a given λ, due to the subsequent reduction of older sibs. However, the frequency of androphilic men among the firstborn individuals is more difficult to predict, and generated data show that it varies with λ, the miscarriage rate, and the interaction between both.

In simulated data, in the absence of miscarriage, the frequency of androphilic men is the same in only-children and in firstborn of larger sibships. This is because, at least in simulated data, later events such as the occurrence of younger sibs have no effect on the sexual orientation of the firstborn. This equality is true despite the fact that these two categories have different sampling probabilities (only-children: *e*^−*λ*^; firstborn of larger sibship: *f*_i_-*e*^−^*^λ^*), because the frequency of androphilic men is calculated within each category. However, in presence of miscarriage, the frequency of androphilic men is higher in only-children than in firstborn of larger sibship. To understand this counter-intuitive phenomenon, let’s consider a situation with two simplified categories: the category “only-children” being composed of real only-children (no sibs miscarried, represented as O), plus children with one older miscarried brother (represented as .O), and the category “firstborn with one younger sib” being composed of real firstborn children with one younger sib (Oo), plus children with an older miscarried brother and one younger sib (.Oo). The frequency of androphilic men is the same in the O and Oo categories, as well as in the .O and .Oo categories. However, the frequency of androphilic men is different in O and .O categories (and in Oo and .Oo), because a miscarried brother, under the MIH, contributes to increase the maternal immune response. Importantly, the sampling probabilities of O and .O are different because their population frequencies are not the same, and similarly the sampling probabilities of Oo and .Oo are different, resulting in different frequency of androphilic men in the O + .O category, compared to the Oo + .Oo category. This reasoning extends to less simplified situation, where the categories “only-children” and “firstborn of larger sibships” are in fact composed of many subcategories having different sampling frequencies. In conclusion, miscarriage generates an “only-child effect”, i.e., a difference in the frequency of androphilic men in only-children, compared to firstborn of larger sibships. Another mechanism has been proposed to explain this “only-child effect”: a specific immune response increasing the odds of same-sex orientation in first pregnancies and reducing subsequent fertility (Blanchard, 2026; Blanchard & Lippa, 2021). Even, if the existence of such additional immune response remains speculative, it could represent another mechanism participating in the “only-child effect”, in addition to miscarriage.

In population samples, the frequency of androphilic men in only-children was overall higher than in firstborn of larger sibship (0.45 *vs* 0.36, respectively, Table S4). This only-child effect suggests the presence of miscarriage (unless another mechanism generates this effect), thus explaining the presence of spurious SBOE when there is an FBOE. The values estimated here to account of the observed SBOE (between 37% and 57%, Table 1) seem higher than the usually reported miscarriage frequency. For example, miscarriage frequency is reported to be around 19% in Iran (Hojati et al., 2025), and around 11% in Canada (Strumpf et al., 2021), although the miscarriage frequency required to explain the SBOE is ∼58% and ∼37% for the Iranian and Canadian samples, respectively. However, these population estimates are relying on clinical values, thus not detecting early pregnancy losses. One key question is determining the developmental stage at which the relevant male-specific antigens are expressed, as a miscarriage before that stage cannot trigger an immune response. The relevant miscarriage frequency should thus be estimated from events after that stage. The Y-linked neuroligin protein NLGN4Y, which has been proposed as a possible candidate for this male-specific antigen in the brain (Blanchard, 2004; Bogaert et al., 2018), is expressed in the nervous system at least 7.5 weeks post-conception (Buonocore et al., 2025), although it may be also expressed in the vascular system (Bottos et al., 2011). In any case, other male-specific antigens than NLGN4Y could be involved, and SRY, which starts sexual differentiation, is first expressed around the 6^th^ weeks post-fertilization (Mamsen et al., 2017). Neural development begins in the third week of gestation, with primary neurulation occurring during weeks 3-4; the brain vesicles then begin to differentiate from the neural tube in the fourth week (Donovan & Cascella, 2026). A minimal conservative miscarriage estimate should thus include early pregnancy losses (defined here as those occurring after blastocyst implantation in the endometrium), and are reported to be ∼30% (Ellish et al., 1996; Wilcox et al., 1988), ∼40% (Chard, 1991; Macklon, 2002), or ∼42-45% (Taylor et al., 2020). Moreover, these rates may be increasing, as the two primary risk factors for pregnancy loss, maternal age and body mass index, have risen significantly in recent decades (Andersen et al., 2000; Genovese & McQueen, 2023; Magnus et al., 2019). Consequently, contemporary miscarriage rates might exceed earlier estimates, complicating direct comparisons with older population samples. Some population estimates are higher than the maximum (45%) reported in the literature (e.g., Czech: 57%; Greece: 58%; Iran: 57%). This possibly suggest that miscarriage rates, even at their highest realistic estimates, may be insufficient to fully explain the observed SBOE if only a spurious effect is assumed, at least in some populations. In other words, mechanisms beyond those proposed by the original maternal immune hypothesis (MIH) may be at play, potentially generating a non-spurious SBOE. However, due to the uncertainty in estimates from the literature, it remains challenging to determine whether the observed SBOE can be fully attributed or not to pregnancy loss alone.

It has previously been suggested that the classical MIH could be extended by considering that the maternal immune response could be triggered primarily by autosomal antigens rather than by specific male antigens (Kabátek & Blanchard, 2024); thus, female pregnancies could also contribute to the maternal immune response and generate a true SBOE alongside the FBOE. However, this hypothesis does not explain why such non-spurious SBOE is not apparent in most studies, particularly those showing a FBOE from populations displaying a low maternal fertility, thus a low spurious SBOE.

However, it is possible that other mechanisms outside the explanatory field of the MIH are operating, and they remain to be identified and evaluated. More generally, birth order impacts traits other than sexual orientation, such as body morphology (e.g., Kwok et al., 2016), life-history traits (e.g., Skjærvø & Røskaft, 2013), personality (Ashton & Lee, 2025), and health (Kramer et al., 2026): whether specific or general explanations are at play remains an interesting issue.

This study has several limitations, although none fundamentally undermine our results. First, only eight population samples were analysed, covering a limited portion of the global geographic diversity. This restricts the generality of the conclusions drawn from the empirical data. Second, some populations had relatively small sample sizes, which may limit the robustness of certain inferences. However, while individual sample values were reported, the primary results are derived from overall analyses across all samples, thereby mitigating the influence of any single sample. Third, our model assumed that miscarriage affects male and female embryos equally, despite possible evidence that male embryos are more susceptible (but see Del Fabro et al., 2011; Ellis & Blanchard, 2001). Nevertheless, even under an extreme scenario where only male embryos were affected by miscarriage, the results remained qualitatively unchanged (data not shown). This suggests that incorporating more realistic differential miscarriage parameters would not alter our conclusions. Finally, the MIH was modelled using four independent parameters: *b*, *slope*, *imr_sd*, and α. Variations in *b* and *slope* were considered, while variations in *imr_sd* could account for the importance of variance in the initial immune response among mothers, and variations in α could reflect the effect of miscarriage on the maternal immune response, respectively. Additionally, not all parameter combinations were explored in the simulations. Finally, several parametrization decisions were arbitrary, such as the linear effect of each son on the maternal immune response, or the logistic effect to assign sexual orientation depending on the level of immune response. Therefore, it remains possible that further phenomena could emerge under different parameter sets or alternative parametrizations of the MIH.

In conclusion, the presence of a FBOE induces a SBOE, driven by the inherent correlation between the number of older brothers and older sisters in population samples. This phenomenon holds true regardless of the underlying mechanism generating the FBOE, including the Maternal Immune Hypothesis (MIH). To mitigate this bias, statistical control for male birth rank, or restricting analyses to a specific birth rank, effectively reduces the occurrence of significant spurious SBOEs to levels consistent with the expected type-I error rate. However, in the presence of miscarriages, the frequency of significant spurious SBOEs may far exceed this baseline. Another consequence of miscarriages is the elevated proportion of androphilic men among only-children, relative to firstborns in larger sibships. Analyses of real-world population data reveal two key findings: first, a significant SBOE persists even after controlling for the FBOE; second, the higher frequency of androphilic men among only-children, compared to firstborns in larger families, strongly suggests a substantial role for miscarriage. Yet, quantitative estimates of the miscarriage rates required to account for the observed SBOE appear unrealistically high compared to the typical miscarriage rates; however, for a more conservative comparison, they appear similar or perhaps higher than rates of early pregnancy losses, perhaps suggesting that a non-spurious SBOE is present, or that additional mechanisms are generating a spurious SBOE. Should the presence of a genuine SBOE be confirmed, its proximal mechanism remains unidentified, presenting a critical avenue for future research.

## Acknowledgements

We thank Ray Blanchard for providing data, and Jakub Fořt, Ray Blanchard, and an anonymous reviewer for useful comments. Computations were performed on the MESO@LR-Platform and ISDM-MESO HPC platforms, funded in the framework of State-region planning contracts (Contrat de plan État-région – CPER) by the French Government, the Occitanie/Pyrénées-Méditerranée Region, Montpellier Méditerranée Métropole, and the University of Montpellier. For the purpose of Open Access, a CC-BY public copyright licence has been applied by the authors to the present document and will be applied to all subsequent versions up to the Author Accepted Manuscript arising from this submission. This is contribution 2026.XXX-SUD of the Institute of Evolutionary Sciences of Montpellier (ISEM, UMR CNRS 5554).

## Funding

The authors declare that they have received no specific funding for this study.

## Conflict of interest disclosure

The authors of this preprint declare that they have no financial conflict of interest with the content of this article. Michel Raymond is a PCI Evol Biol recommender.

## Data, scripts, code, and supplementary information availability

Scripts are available at: https://doi.org/10.5281/zenodo.18879331, and data are available at: https://doi.org/10.5281/zenodo.18910601

## Declaration of AI use

During the preparation of this work, the authors used Emmy in order to improve grammar and style. After using this tool, the authors reviewed and edited the content as needed and take full responsibility for the content of the publication.

## Notes

https://doi.org/10.5281/zenodo.18879331

https://doi.org/10.5281/zenodo.18910601

### Competing Interest Statement

The authors have declared no competing interest.

### Summary of Updates

Minor changes (typo corrections; 3 sentences edited; two references added).

