## Supplementary materials for "The role of miscarriage and sororal birth order in male same-sex orientation: Theoretical predictions and empirical data"

Michel Raymond[1], Ana Aguerre[1], Valerie Durand[1], Menelaos Apostolou[2], Julien Barthes[1], Sarah Nila[3,4], Bambang Suryobroto[3], Mostafa Sadr-Bazzaz[5], Paul L. Vasey[5], Daniel Turek[6], and Pierre-André Crochet[7]

1. ISEM, Univ Montpellier, CNRS, EPHE, IRD, Montpellier, France
2. Department of Social Sciences, University of Nicosia, Cyprus
3. Department of Biology, Faculty of Mathematics and Natural Sciences, IPB University (Bogor Agricultural University), Indonesia
4. Department of Political Science, University College London, London, UK
5. Department of Neuroscience, University of Lethbridge, Lethbridge, Canada
6. Department of Mathematical Sciences, Lafayette College, Easton, PA, USA
7. CEFE, CNRS, Univ Montpellier, EPHE, IRD, Montpellier, France

### Supplementary materials

**Supplementary tables**

**Table S1**. Functions used to model an older sister effect. x = number of older sisters. The type of curve (Family), and the corresponding parameters are indicated.

| Name | Form | Family | Parameters |
|---|---|---|---|
| f1 | $a \cdot \left(1-e^{-bx}\right)$ | Saturating exponential | a, b |
| f2 | $a \cdot \left(1-c \cdot e^{-bx}\right)$ | Saturating exponential | a, b,c |
| f3 | $a+b \cdot x^{\gamma}$ | Geometric | a, b, γ |
| f4 | $\frac{1}{1+e^{-(a+b \cdot x)}}$ | Logistic | a, b |
| f5 | $\frac{1}{1+e^{-(a+b \cdot x+c \cdot x^2)}}$ | Logistic | a, b, c |
| f6 | $\frac{1}{1+e^{-(a+b \cdot x+c \cdot x^2+d \cdot x^3)}}$ | Logistic | a, b, c, d |
| f7 | $\frac{1}{1+e^{-(a+b \cdot x+c \cdot x^2+d \cdot x^3+e \cdot x^4)}}$ | Logistic | a, b, c, d, e |

**Table S2.** WAIC comparisons of different functions to describe the correlative SBOE generated by MIH. Increasing sample size (rows) allows studying the shape of SBOE over a larger range of number of older sisters (*os* range), as categories below 50 individuals are not considered. For a given range of *os*, the sample size *N* required (containing an equal proportion of androphilic and gynephilic men) is a function of the mean population fertility λ. For each line, the lowest WAIC is underlined, and WAIC values lower than the minimum value plus 2 are in bold.

| N | | *os* range | Function | | | | | | |
|---|---|---|---|---|---|---|---|---|---|
| | | | Saturating exponential | | Geometric | Logistic | | | |
| $\lambda = 3$ | $\lambda = 5$ | | f1 | f2 | f3 | f4 | f5 | f6 | f7 |
| 1,200 | 600 | 0 – 3 | **731.2** | 737.2 | **731.0** | <u>**730.9**</u> | **731.8** | **732.9** | **732.5** |
| 10,200 | 3,000 | 0 – 5 | 3989.7 | **3983.4** | **3984.1** | 3987.6 | <u>**3982.7**</u> | **3983.5** | 3985.0 |
| > 60,000 | 6,600 | 0 – 6 | <u>**8889.6**</u> | **8891.3** | 8903.8 | 8928.8 | 8900.2 | 8892.0 | 8892.5 |

Table S3

**Table S3.** Demography parameters of the population samples. The number of androphilic (Andro) and gynephilic (Gyne) men are indicated, as well as the correlation between the number of older brothers and older sisters ($\tau$) and the associated *P*-value, the mean fertility calculated from the whole sample (all), or only from the gynephilic men (gyn.), and the under/over-dispersion parameter ($v$) describing the variance ($\lambda/v$) of the distribution ($v$ = 1 for a standard Poisson distribution of parameter $\lambda$), with their associated SEM values. “All” refers to the sum for all samples (for N), to the mean over all sample values (for τ), or to the value over the overall dataset (for fertility). Significant *P*-values ($P < 0.05$) are in bold.

| | N | | Correlation (ob,os) | | Fertility | | | | | |
|---|---|---|---|---|---|---|---|---|---|---|
| Sample | Andro | Gyne | τ | *P*-value | Mean (all) | (SEM) | Mean (gyn.) | (SEM) | *v* | (SEM) |
| Canada | 302 | 434 | 0.204 | **$1.7\ 10^{-9}$** | 2.326 | (0.056) | 2.251 | (0.072) | 0.75 | (0.06) |
| Czech | 698 | 843 | 0.050 | **0.039** | 1.388 | (0.030) | 1.415 | (0.041) | 1.46 | (0.07) |
| France | 253 | 241 | 0.007 | 0.864 | 1.623 | (0.057) | 1.680 | (0.084) | 1.28 | (0.11) |
| Greece | 183 | 593 | 0.072 | **0.036** | 1.465 | (0.043) | 1.433 | (0.049) | 1.19 | (0.09) |
| Indonesia | 79 | 62 | 0.245 | **0.001** | 3.596 | (0.160) | 3.387 | (0.234) | 0.34 | (0.08) |
| Iran | 420 | 239 | 0.274 | **$<10^{-10}$** | 2.023 | (0.055) | 1.912 | (0.089) | 0.83 | (0.07) |
| Polynesia | 149 | 231 | 0.385 | **$<10^{-10}$** | 3.632 | (0.098) | 3.710 | (0.127) | 0.45 | (0.05) |
| Samoa | 358 | 703 | 0.354 | **$<10^{-10}$** | 4.994 | (0.069) | 4.663 | (0.081) | 0.79 | (0.04) |
| All | 2476 | 3346 | 0.199 | **0.008** | 2.472 | (0.021) | 1.459 | (0.021) | - | - |

Table S4

**Table S4.** Frequency of androphilic men in only-children and firstborn with younger siblings. The number of individuals in each category is given, the frequency of androphilic men ($f_H$), and the odd-ratio and *P*-value for the corresponding 2×2 contingency table, for each population sample. “All” refers to the sum of the individual numbers, the frequency of androphilic men over the whole dataset, or to the combined one-tailed *P*-values using Fisher’s method. Significant *P*-values ($P < 0.05$) are in bold.

| | Only-children | | | First-born | | | 2x2 contingency table | |
|---|---|---|---|---|---|---|---|---|
| Sample | Homo | Hetero | $f_H$ | Homo | Hetero | $f_H$ | Odd-ratio | *P*-value |
| Canada | 24 | 28 | 0.462 | 94 | 181 | 0.342 | 0.607 | 0.069 |
| Czech | 104 | 120 | 0.464 | 249 | 382 | 0.395 | 0.752 | **0.041** |
| France | 36 | 29 | 0.554 | 67 | 92 | 0.421 | 0.588 | **0.049** |
| Greece | 19 | 86 | 0.181 | 52 | 250 | 0.172 | 0.942 | 0.472 |
| Indonesia | 1 | 3 | 0.250 | 13 | 23 | 0.361 | 1.675 | 0.836 |
| Iran | 45 | 19 | 0.703 | 119 | 79 | 0.601 | 0.637 | 0.093 |
| Polynesia | 8 | 6 | 0.571 | 40 | 70 | 0.364 | 0.432 | 0.114 |
| Samoa | 9 | 9 | 0.500 | 40 | 142 | 0.220 | 0.284 | **0.013** |
| All | 246 | 300 | 0.451 | 674 | 1219 | 0.356 | - | **0.002** |

**Table S5.** Younger sibling effects from population data. A binomial regression was performed with the sexual orientation as the response variables, the number of younger brothers (or younger sisters), the number of older brothers, and the number of sibs as independent variables. The slope (β) of younger brothers, or younger sisters, is given, with their corresponding SEM and *P*-value, for each sample. 'All' refers to values generated when all population samples were pooled. Significant ($P < 0.05$) *P*-values are in bold.

| | Younger brother | | | Younger sister | | |
|---|---|---|---|---|---|---|
| Sample | β | (SEM) | P-value | β | (SEM) | P-value |
| Canada | -0.166 | (0.115) | 0.1462 | -0.017 | (0.119) | 0.8835 |
| Czech | -0.003 | (0.096) | 0.979 | -0.248 | (0.104) | **0.0164** |
| Greece | -0.201 | (0.157) | 0.1942 | -0.025 | (0.173) | 0.8854 |
| Indonesia | 0.065 | (0.223) | 0.7688 | -0.088 | (0.215) | 0.6823 |
| Iran | -0.28 | (0.151) | 0.0642 | -0.136 | (0.151) | 0.3708 |
| Polynesia | 0.057 | (0.126) | 0.6493 | -0.057 | (0.132) | 0.6647 |
| Samoa | -0.127 | (0.073) | 0.0817 | -0.158 | (0.072) | **0.0262** |
| All | -0.113 | (0.041) | **0.0063** | -0.168 | (0.043) | **1e-04** |

**Supplementary figures**

**Figure S1.** Estimating the frequency of miscarriage from the slope of SBOE. Population samples were generated with a known frequency of miscarriage ("True *fm*"), and this frequency was estimated ("Estimated *fm*") from the value of the slope of SBOE. The circles represent the mean of at least 200 replicates (dots) for each value of *fm*. The dotted line represents the expected curve when the estimated *fm* equals the true *fm* value.

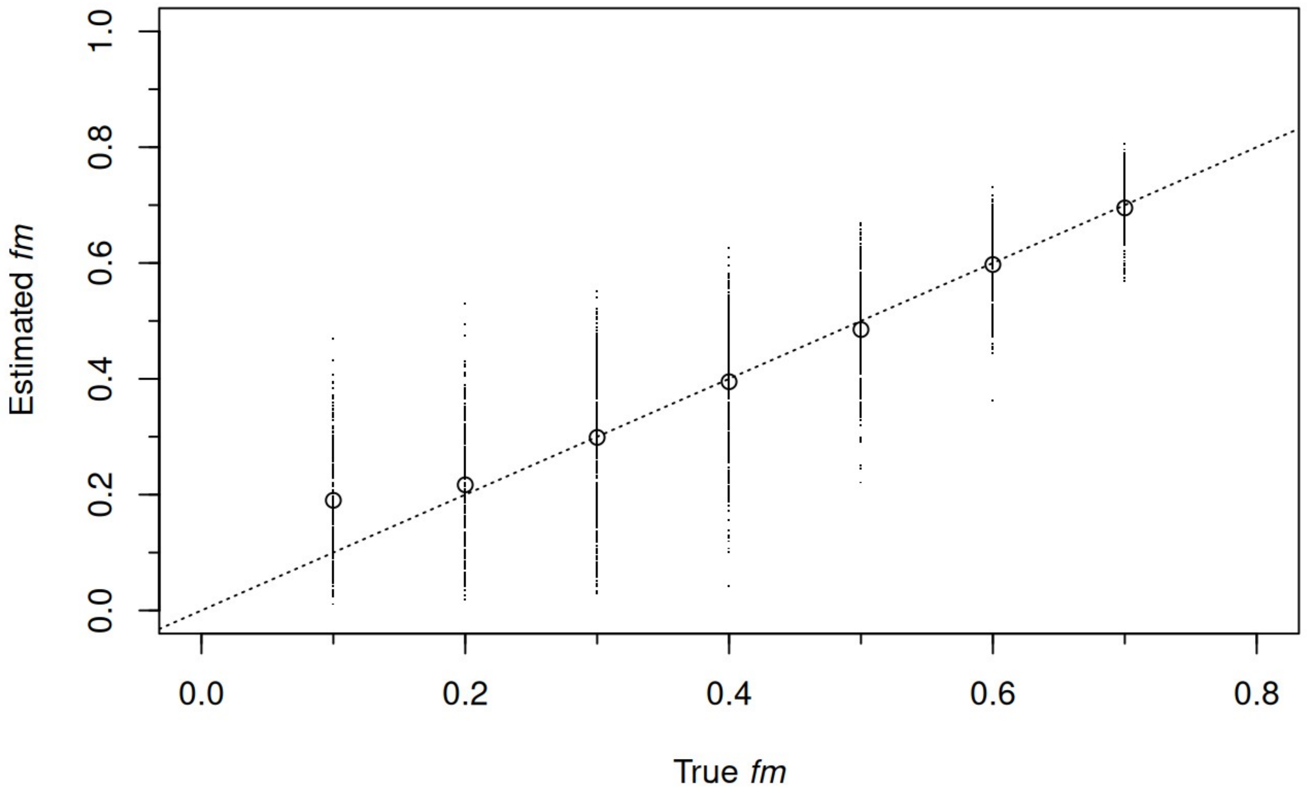

**Figure S2**. Effect of miscarriage frequency on the frequency of firstborn (including only-children) androphilic men. Data were generated with the same number of androphilic or gynephilic men, and a total size of N = 2,400, a miscarriage cost $\alpha = 25\%$. A. The mean population fecundity is $\lambda_A = 5$, thus the observed fertility is $\lambda_A(1- fm)$, with *fm* being the miscarriage frequency. B. The mean population fecundity is $\lambda_B = \lambda_A/(1- fm)$, so that the observed mean fertility is $\lambda_A$. The circles represent the mean of each replicate (N > 50, individual dots), for each frequency of miscarriage.

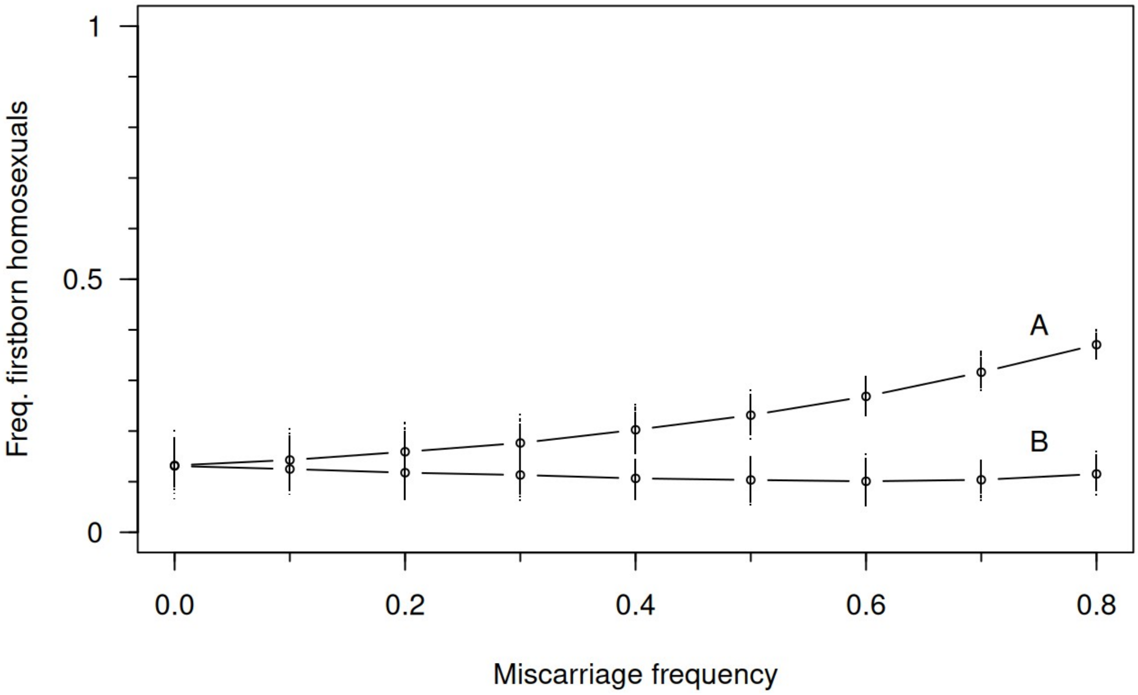

**Figure S3.** Effect of the amplification parameter (*n*) on the distribution estimates of *fm* from the slope of SBOE. The distribution of the *fm* estimates is shown for each population sample and for all samples (“All”), and for *n* = 1 (grey), *n* = 10 (red), and *n* = 50 (blue). The mean of each distribution is indicated by a dot below the histograms, with the same colour code.

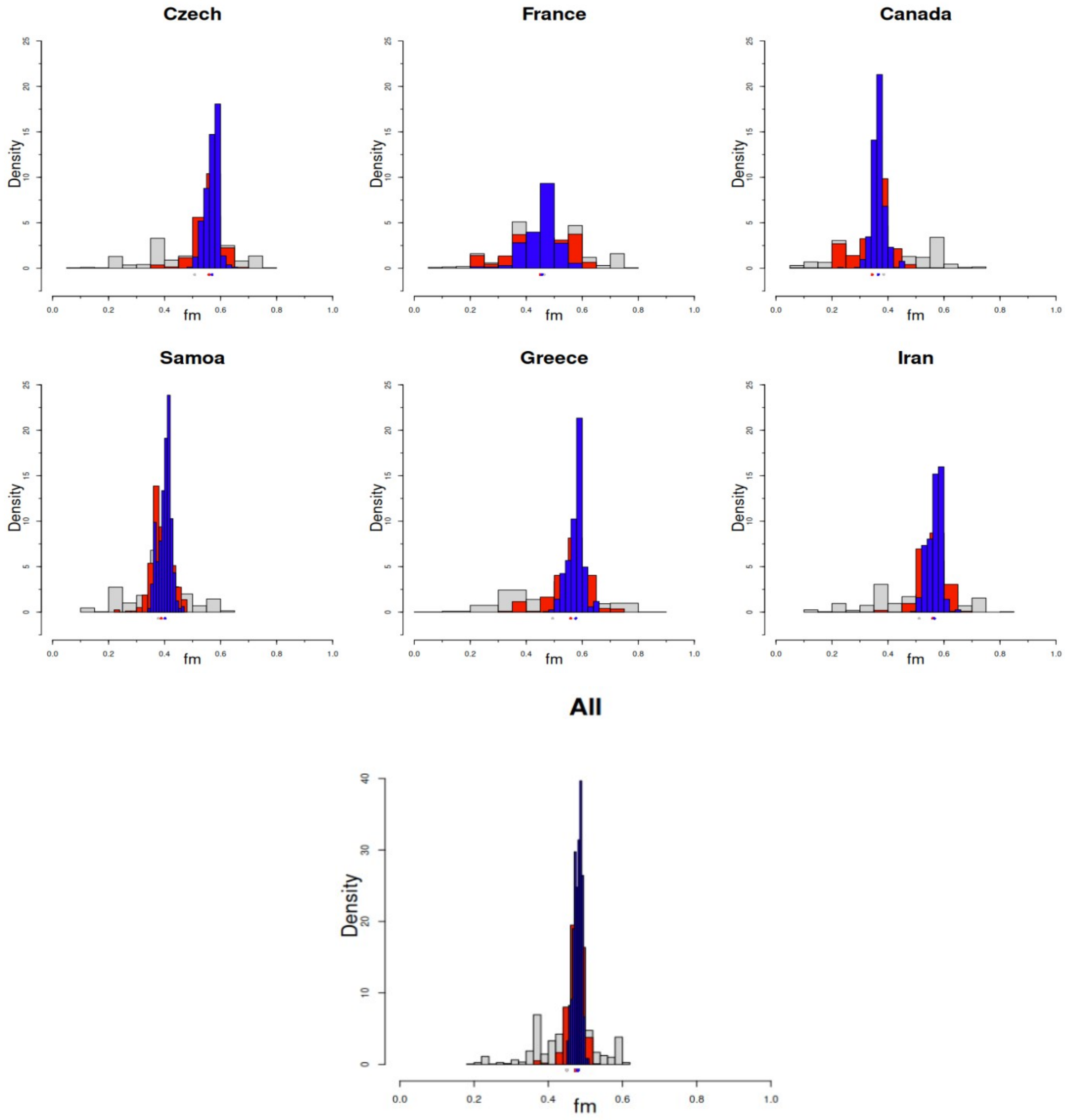

**Figure S4.** Effect of miscarriage frequency on type-I error for detecting an SBOE. Data were generated under the MIH for various values of miscarriage affecting only male embryos, and SBOE was detected using a logistic regression, controlling for the number of lived older brothers. Each point depicts the proportion of significant SBOE over 400 independent replicate samples, with the corresponding 95% confidence interval. Each sample is composed of the same number of androphilic or gynephilic men, with the total size being N = 600 (empty circles), N = 1,200 (empty squares), or N = 2,400 (empty rhombus). The dotted horizontal line indicates the false rejection rate of 2.5%.

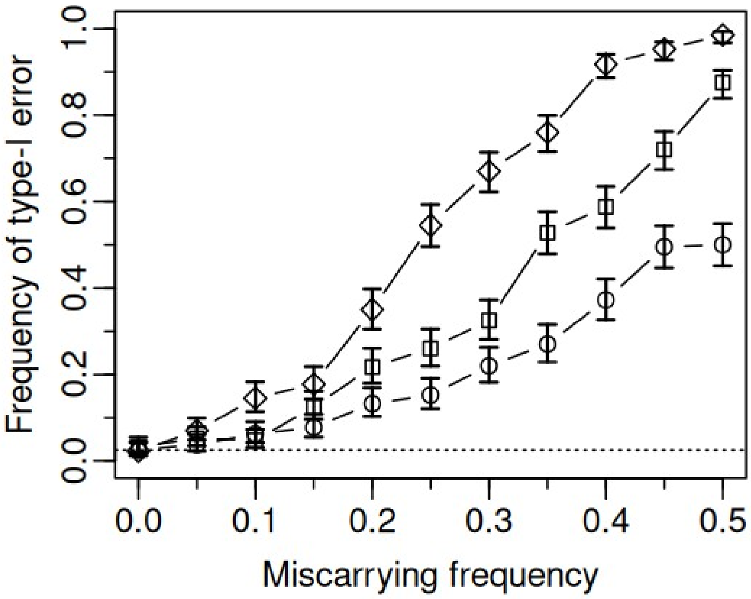

**Figure S5.** Effect of miscarriage frequency on type-I error for detecting an opposite younger sib effect. Data were generated under the MIH for various values of miscarriage affecting both male and female embryos, and the younger brother effect was detected using a logistic regression, controlling for the number of lived older brothers. Each point depicts the proportion of significant younger brother effect with a negative slope, over 400 independent replicate samples, with the corresponding 95% confidence interval. Each sample is composed of the same number of androphilic or gynephilic men, with the total size being N = 600 (empty circles), N = 1,200 (empty squares), or N = 2,400 (empty rhombus). The proportion of significant younger brother effect when controlling for the initial number of older brother (before miscarriage) is depicted by full small rhombus (only N = 2,400 is shown). The dotted horizontal line indicates the false rejection rate of 2.5%.

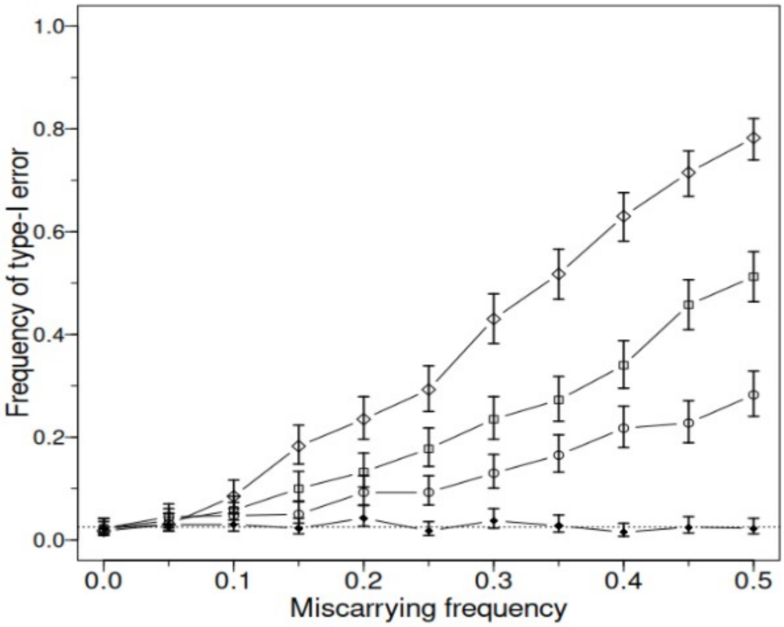
